# gsQTL: Associating genetic risk variants with gene sets by exploiting their shared variability

**DOI:** 10.1101/2024.09.13.612853

**Authors:** Gerard A. Bouland, Niccolò Tesi, Ahmed Mahfouz, Marcel J.T. Reinders

## Abstract

To investigate the functional significance of genetic risk loci identified through genome-wide association studies (GWASs), genetic loci are linked to genes based on their capacity to account for variation in gene expression, resulting in expression quantitative trait loci (eQTL). Following this, gene set analyses are commonly used to gain insights into functionality. However, the efficacy of this approach is hampered by small effect sizes and the burden of multiple testing. We propose an alternative approach: instead of examining the cumulative associations of individual genes within a gene set, we consider the collective variation of the entire gene set. We introduce the concept of gene set QTL (gsQTL), and show it to be more adept at identifying links between genetic risk variants and specific gene sets. Notably, gsQTL experiences less susceptibility to inflation or deflation of significant enrichments compared with conventional methods. Furthermore, we demonstrate the broader applicability of shared variability within gene sets. This is evident in scenarios such as the coordinated regulation of genes by a transcription factor or coordinated differential expression.

## Background

Genome-wide association studies (GWASs) identify genomic loci linked to specific traits, but their impact is hard to understand as the majority of associated loci fall in non-coding and intergenic regions of the genome^1,2^. To determine the functional significance of genetic variants, they are often linked to changes in mRNA expression in bulk RNAseq data (eQTLs)^3^ or single cell RNAseq data (sc-eQTLs)^4–7^, as well as other molecular data, such as lipids (lipid-QTLs^8^), metabolites (mQTLs^9,10^), and microRNA expression (miQTLs^11^). Once these QTLs are identified, gene set analysis is commonly used to identify affected pathways^12–16^. In the context of complex diseases, this approach typically begins with multiple SNPs, as these are associated with multiple genes, enabling overrepresentation analysis (ORA). However, when focusing on a single variant, this becomes challenging due to the limited number of significantly associated genes, which arises from the burden of multiple testing and the small effect sizes observed in QTL analyses^3,17^, especially when associations between a single variant and the whole transcriptome is considered (trans-eQTLs). Another approach is gene set enrichment analysis (GSEA), which in contrast to ORA considers the entire list of genes ranked by their change in expression levels or other relevant metrics (e.g., p-value, effect size) without requiring a predefined threshold for differential expression. It assesses whether a predefined gene set shows significant, consistent differences in expression across the entire ranked list. However, GSEA approaches often yield higher numbers of significantly enriched gene sets compared to ORA approaches, but also have been found to have elevated false positive rates^18^. To address the limitations of functional enrichment analysis of GWAS loci, we propose directly assessing the impact of SNPs on the overall variability of expression of a gene set, contrasting with traditional post-hoc aggregation approaches (e.g. ORA or GSEA). Our gene set QTL (gsQTL) approach starts by combining genes into gene sets based on known interactions (gene sets). Previous studies have demonstrated the superiority of principal component analysis (PCA) over other popular methods for hidden variable inference for QTL related analyses^19^. However, principal components frequently capture gene expression variance that arises from a mix of various biological and technical sources within a single component^20^. Our approach guides the identification of principal components by calculating them within a highly controlled environment, specifically focusing on predefined gene sets. This approach effectively isolates the biological variance directly associated with the specific biological factor represented by the gene set. Furthermore, unlike other methodologies^21^, our components are immediately interpretable due to the inclusion of the biological factor (gene set) that directly links the corresponding genes.

The advantages of this approach for QTL analyses include the ability to capture collective variation within a gene set using PCA, which can lead to stronger associations with risk variants, even when the effect size of the individual genes are modest, as demonstrated in **Fig. 1**. Moreover, the number of gene sets is lower than the number of genes, which reduces the burden of multiple testing correction, and hence increases statistical power. In addition, multiple genes are tested at once for each SNP and these genes do not need to be located near the SNP, allowing for an integration of cis and trans regulatory effects.

**Figure 1:**
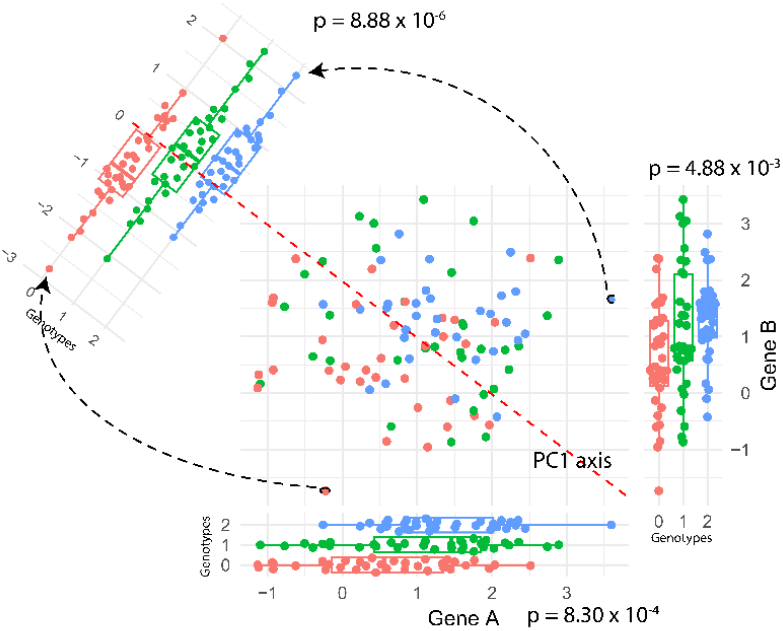
Graphical explanation with simulated data of the proposed gene set QTL (gsQTL) approach. Illustrated are the effects of three genotypes of a SNP on the expression of Gene A (x-axis) and Gene B (y-axis) that are together forming one gene set, across a set of measured samples. When we inspect the expression of an individual gene for both variations (box plots at bottom and right side of the figure), there is a significant difference in expression, however, by representing the shared variation of the two genes within the gene set, using the first component of a principle component analysis (PCA), we can observe that the shared variation in the gene set shows a much stronger significant difference in expression when the SNP varies (box plot at diagonal of figure).

## Results

Using six jointly pre-processed Alzheimer’s Disease (AD) scRNAseq datasets^23–28^, including genetics data from 666 individuals (N=249 AD cases, N=242 healthy controls, and N = 175 with mild cognitive impairment with other causes than AD), we demonstrate that gsQTL detects novel functional implications of AD-related SNPs and indeed has more power to detect them than current post-hoc approaches. gsQTL analyses were performed on previously identified AD risk variants^29,30^. Following pre-processing, 44 variants were retained for testing. For seven major brain cell types (excitatory neurons, inhibitory neurons, astrocytes, oligodendrocytes, microglia, OPCs and endothelial cells), we report 66 significant gsQTLs comprised of 30 AD risk variants and 59 unique gene sets, including microRNA targeted genes^31^, metabolite interacting genes^32^, and KEGG pathway-related gene sets^33^ (**Fig. 2a** , **Sup. Tables 1-3**).

**Figure 2:**
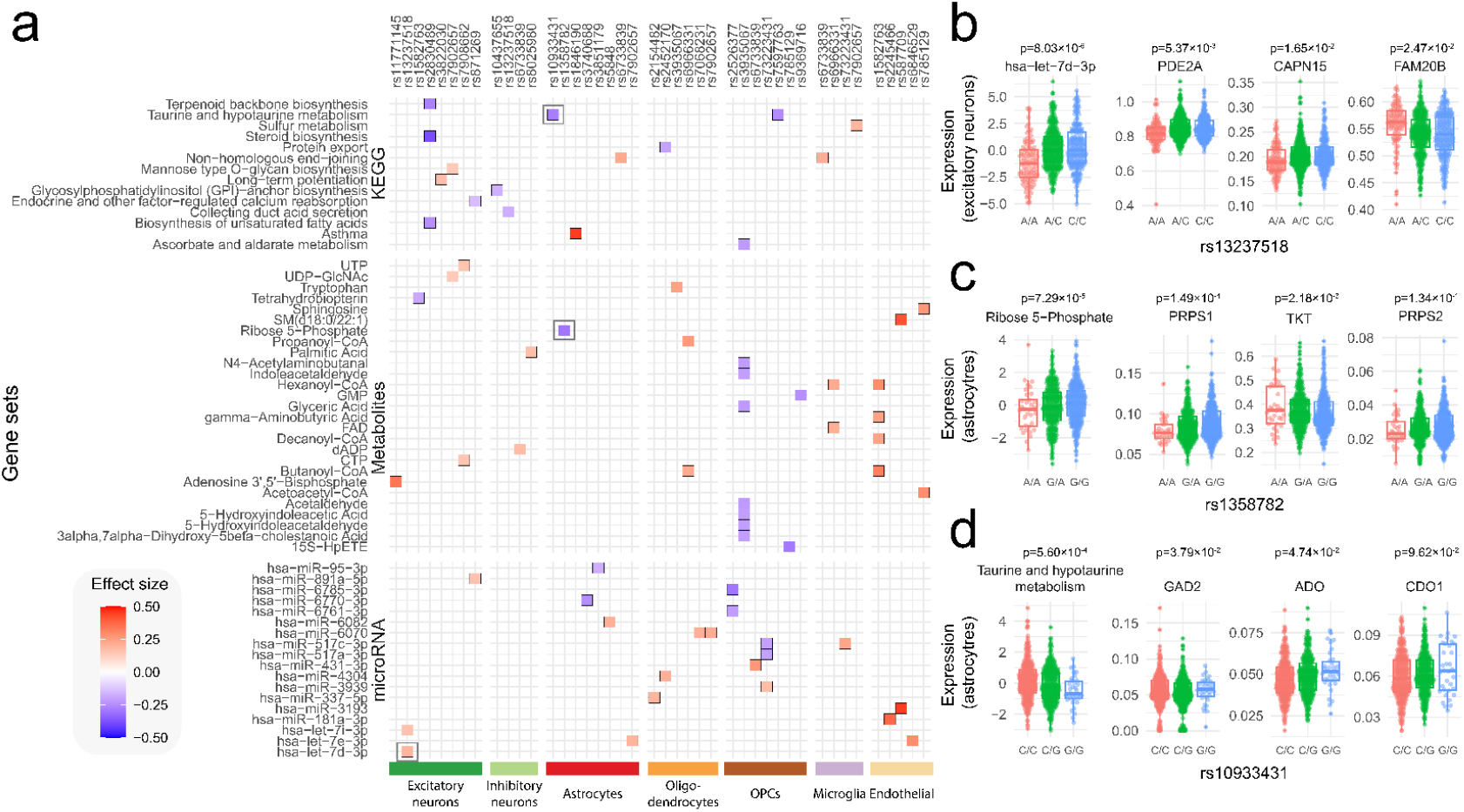
**a**, Significant association between gene sets (rows) and genetic variant (columns) grouped by major cell type (bottom colors) and with the effect size (β, red-blue gradient) of the association between gene set and risk variant. **b**,**c**,**d**, Boxplots of a selected set of gsQTL: **b**, hsa-let-7d-3p-rs13237518, **c** Ribose 5-Phosphate-rs1358782 and **d**, taurine and hypotaurine metabolism-rs10933431, with the genotypes (x-axis) of the respective SNPs and the expression levels (y-axis) of the whole gene set as well as three individual genes that belong to the respective gene sets.

For 53 of the 66 significant gsQTLs, the association of the gene set with the variant had a smaller nominal p-value than the association between the variant and any of the individual genes within the respective gene set (**Supp. Fig 1, Supp Table 4**), showing empirically that the shared variation among genes within a gene set can indeed yield stronger associations with a variant compared to those observed when considering only individual genes of that gene set. Among the significant microRNA gene sets identified, hsa-let-7d-3p (p=8.03×10^-6^, **Fig. 2b**) and hsa-let-7i-3p (p=2.36×10^-4^) showed significant associations with rs13237518 (intronic of *TMEM106B*) in excitatory neurons, both of which have been previously implicated in AD pathology^34,35^. Additionally, we identified hsa-miR-6761-3p (5.90×10^-4^) associated with rs2526377 (intronic of *TSPOAP1*) in OPCs, also a microRNA previously implicated in AD pathology^36^. Further, through the gsQTL analysis considering the gene sets interacting with metabolites, we uncover a potential role of rs1358782 (intronic of *RBCK1*) in pentose phosphate pathway, through the gsQTL in astrocytes with Ribose 5-Phosphate^37^ (p=7.29×10^-5^, **Fig. 2c**). And, among the KEGG pathways, we identified a significant association between taurine and hypotaurine metabolism (p=5.60×10^-4^, **Fig. 2d**) and rs10933431 (intronic of *INPP5D*) in astrocytes. This finding is particularly interesting given the growing recognition of astrocytes’ crucial role in cognitive functioning^38,39^ and the neuroprotective properties of taurine, which is believed to enhance cognitive performance^40^. These results suggest that the AD-risk allele of rs10933431 might be involved in a dysregulation of taurine and hypotaurine metabolism, specifically in astrocytes. Collectively, these comparisons and results demonstrate the capacity of gsQTLs to reveal associations with gene sets that may be overlooked when focusing solely on individual genes initially.

To compare gsQTLs with more traditional post-hoc approaches, we considered: 1) overrepresentation analysis (ORA) using the fisher exact test, and 2) gene set enrichment analysis (GSEA) using the R-package fgsea^41^. With ORA, we identified only two gene sets significantly associated with a variant (**Supp. Table5-7**), which is consistent with the limitations of this approach, which is generally more effective when multiple SNPs are analyzed simultaneously. With GSEA, we detected 246 gene sets significantly associated with a variant, including 188 KEGG pathways, 46 metabolite interacting gene sets and 30 microRNA targeted gene sets (**Supp. Table8-10**), comprising 43 unique variants and 81 unique gene sets. Only 5 gene sets-variants-cell type combination overlapped with those identified by our gsQTL method (**Fig. 3a**). Upon closer examination, we observed that the ribosome pathway was linked to 28 distinct variants specifically in excitatory neurons. Despite this, the ribosome pathway did not emerge as significant in our gsQTL analysis, even though 111 out of 127 ribosomal genes showed moderate associations (p≤0.05) with at least one variant. These ribosomal genes exhibit a high degree of collinearity amongst each other, resulting in the shared variability being captured predominantly by the first principal component in PCA (**Fig. 3b**). But this signal is not strong enough to find an association with any of the variants. As GSEA has been reported to have elevated false positive rates^18^, this might be the case here too, suggesting that our gsQTL analysis effectively accounts for the inflation of association signals caused by collinearity among genes.

**Figure 3:**
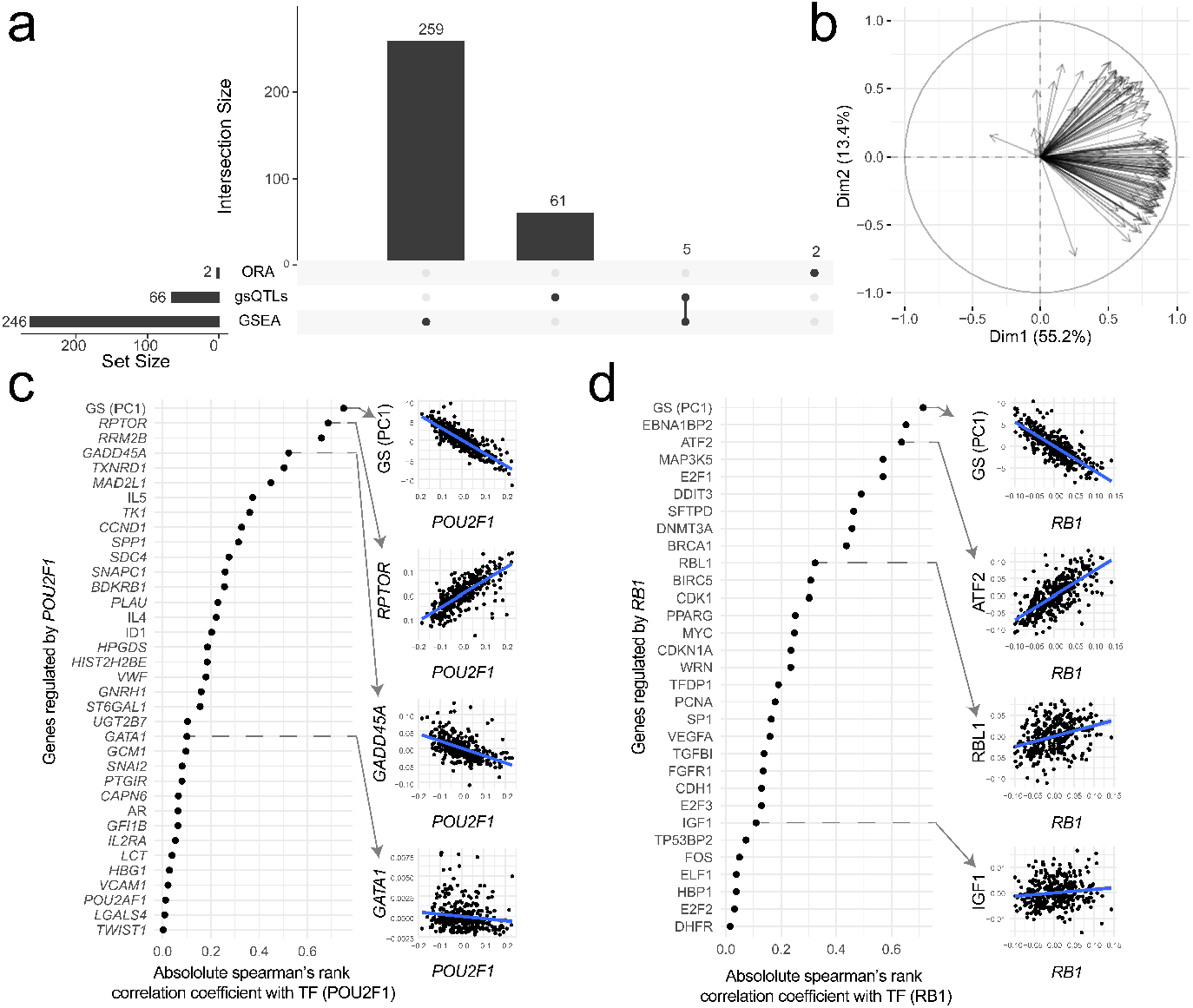
**a**, Overlap between the gene sets identified with ORA, gsQTLs and GSEA. **b**, PCA correlation circle plot of all genes comprising the ribosome KEGG pathway and expressed in our excitatory neuron dataset. The vectors represent the individual ribosomal genes. The closer the vector to each other, the more similar the correlations of the respective genes with respect to the PCs are. **c**,**d** Absolute correlations between the **c**, TF POU2F1 and **d**, TF RB1 and their targets (x-axis) and the gene set as a whole (GS PC1), with each of the individual targets and the gene set as a whole represented on the y-axis. On the right of the respective plots scatter plots showing the TF expression(x-axis) and target expressions (y-axis) in excitatory neurons where every dot represents a donor.

In our approach, we assume that shared variability among genes reflects a common underlying regulatory mechanism, such as a microRNA targeting a gene set or participation in a KEGG pathway. By analyzing the association between expression changes in TFs and the coordinated expression shifts in their downstream targets, we aimed to test how broadly our shared variation approach could be applied across different regulatory contexts. More specifically, we computed the correlation between the expression of the TF and the shared representation of the set of genes targeted by that TF. Using data from all individuals (N = 706) in the excitatory neuron dataset and regressing out potential confounding variables (age, sex, dataset and braak stage), we conducted the gsQTL analysis for 256 TFs obtained from the TRUST transcription factor database^42^ that overlapped with the measured expression data. We observed that for 42 TFs (16%), their expression patterns correlated with the shared representation of their target genes (P≤ 4.98×10^-46^, r ≥ 0.50). Interestingly, for 43 of the TFs, the shared representation showed a stronger correlation with the TF than any of the individual TF target genes (**Fig. 3c-d**), underscoring the considerable value of shared maximum variability within a gene set in association analyses.

To further explore the utility of our shared variation approach, we investigated whether a shared representation of a gene set is more effective in detecting differential expression across conditions compared to traditional methods involving differential expression per gene followed by ORA or GSEA for functional interpretation. For this purpose, we utilized an acute myeloid leukaemia (AML) bulk RNAseq dataset comprising 3,383 individuals^43^. Our aim was to identify metabolic gene sets exhibiting differential expression between AML-subtypes characterized by genetic abnormalities: KMT2A.1 (N_individuals_ = 540), RUNX1_RUNX1T1 (N_individuals_ = 300), FLT3_ITD (N_individuals_ = 661), CBFB_MYH11 (N_individuals_ = 310) and NPM1 (N_individuals_ = 508), TET2 (N_individuals_ = 194). We evaluated 233 metabolome-related gene sets and tested whether their shared representation was differentially expressed between individuals with and without the respective genetic abnormalities. Metabolic gene sets were significantly (PFDR ≤ 0.05) differentially expressed in FLT3_ITD (N_gene sets_ = 40), followed by KMT2A.1 (N_gene sets_ = 39), CBFB_MYH11 (N_gene sets_ = 32), RUNX1_RUNX1T1 (N_gene sets_ = 31) and NPM1 (N_gene sets_ = 27). Moreover, most metabolic gene sets were differentially expressed in more than two subtypes (**Supp. Fig. 3a**). For example, the most pronounced differential expression between blood and bone marrow was observed for the metabolic gene set related to *phosphatidylcholine* in the KMT2A.1 subtype (P_FDR_ = 3.77 × 10^-53^, **Supp. Fig. 3b**), which was not detected using the post-hoc analysis approach. Notably, phosphatidylcholine is the product of a reaction catalyzed by genes of the LPC acyltransferase family. *LPCAT1*, a member of this family, is suggested as biomarker to guide treatment choice in AML patients.^44^ When applying the ORA approach, only five metabolic gene sets are significantly associated with an AML-subtype, whereas with GSEA, no metabolic gene sets are significantly associated with all AML-subtypes, showing that our shared variability approach identifies associations of genetic AML abnormalities with gene sets that would otherwise go unnoticed.

## Conclusion

We present an approach for exploring the functional implications of genetic risk variants on gene expression, biomolecules, and pathways. Rather than focusing on individual gene associations, our method identifies and examines the shared variability of a gene set in relation to genetic risk variants. By defining gene sets based on interactions with various biomolecules such as metabolites or microRNAs, but also based on biological pathways, our approach provides additional context to cell type specific changes associated with genetic risk variants. Our findings underscore that a priori identification of shared variability within gene sets, via PCA, facilitates the discovery of putative coordinated expression changes. These findings can be broadly applied, as demonstrated in our study, for associating with genetic risk variants, coordinating the regulation of genes targeted by a transcription factor, or coordinated differential expression.

## Availability of data and materials

All codes and analysis results in this paper are publicly available at GitHub: https://github.com/gbouland/gsQTLs. An R-package of gsQTLs is available at GitHub: https://github.com/gbouland/gsQTL The single-cell RNAseq, genetics and clinical datasets are available on AD Knowledge Portal and are available under the following accession codes: syn16780177, syn18485175, syn31512863, syn52293433, syn3157322, syn17008939 and syn28257618. The AML data is available at https://osf.io/wq7gx/.

## Contributions

GAB, AM, and MJTR conceived the study designed the experiments. GAB performed all experiments and drafted the manuscript. NT pre-processed and combined the genetics data. GAB, NT, AM, and MJTR reviewed and approved the manuscript.

## Online methods

### Single-cell RNAseq data

Five 10x single-cell RNAseq (scRNAseq) datasets were acquired from AMP-AD knowledge portal of which the subjects were participants of the Religious Orders Study and the Memory and Aging Project (ROS/MAP)^45^. The Seattle Alzheimer’s Disease Brain Cell Atlas (SEA-AD) was obtained from (https://registry.opendata.aws/allen-sea-ad-atlas/). The first dataset (**BA9**, ID: syn16780177) consisted of 24 subjects and originated from the dorsolateral prefrontal cortex (DLPFC), specifically Brodmann area 9 (BA9). Raw fastq files were obtained of this dataset. The second dataset (***BA10***, ID: syn18485175) consisted of 48 subjects and originated from the prefrontal cortex (PFC), specifically BA10. A count matrix was obtained of this dataset as it was already processed with CellRanger aligning reads to the hg38 genome^25^. The third dataset (**TREM**, ID: syn18485175) consisted of 32 subjects and originated from the DLPFC, BA9 and BA46. Of this dataset a count matrix was obtained as it was also processed with CellRanger aligning reads to the hg38 genome^27^. The fourth dataset (**DLPFC2**: ID: syn31512863) consisted of 424 individuals. Of this dataset also a count matrix was obtained as it was also processed with CellRanger aligning reads to the hg38 genome^24^. The fifth dataset (**MIT:** ID: syn52293433) consisted of 427 individuals. Of this dataset also a count matrix was obtained as it was also processed with CellRanger aligning reads to the hg38 genome^28^. The sixth dataset (**SEA-AD**) consisted of 89 middle temporal gyrus samples, 23 of which were diagnosed with AD and 32 were specified as CT^23^. Of this dataset the raw count matrix was acquired.

### Clinical data

Clinical data were acquired from the AMP-AD knowledge portal (ID: syn3157322). The variable cogdx was used to characterize controls (CT), Alzheimer’s disease (AD) and other (O). Cogdx represents the clinical consensus diagnosis of cognitive status at time of death and is indicated with a value ranging from one to six. A value of one represents no cognitive impairment (CI), as such, individuals with a cogdx of one were characterized as CT. A value of four represents Alzheimer’s dementia and no other cause of CI, as such, these individuals were characterized as AD. The remaining values represent mild CI and/or other causes for dementia and these individuals were characterized as O. Besides clinical diagnosis, *APOE* genotype, Braak stage, sex, and age at time of death was also available. However, age at time of death is censored above the age 90 years. Of the **SEA-AD** datasets the clinical data were acquired from the corresponding source. For both datasets; age, sex, clinical diagnosis and Braak stage were available.

### Genetics data

Genotyping data were sourced from the Synapse AD portal and consisted of 3 batches. Batch 1 and batch 2 (SynID: syn17008939) included 1709 and 382 individuals, respectively, from the ROSMAP study^45^. Batch 3 (SynID: syn28257618) included 95 samples from SEA-AD study^23^. Batch 2 and batch 3 data were aligned to GRCh37 (hg19), while batch 1 data was aligned to GRCh36 (hg18) and lifted over to GRCh37 (hg19). Standard quality control was applied to each batch independently (variant call rate >98%, individual call rate >98%, and deviation from Hardy-Weinberg was considered significant at p<1e^-6^). Variant ID, strand, and allele frequencies were compared, for each batch, to the Haplotype Reference Consortium (HRC, HRC-1000G-check-bim-v4.2.7.pl)^46^. Genotyping data were combined and high-quality genotyping was ensured (variant call rate >98%, individual call rate >98%). All autosomal variants were submitted to the TOPMED imputation server (https://imputation.biodatacatalyst.nhlbi.nih.gov). The server uses Eagle (v2.4) to phase data and imputation to the reference panel (TOPMED R2 v1.0) was performed with Minimac4^47–49^. A total of 2,115 individuals passed quality control. Prior to analysis, we extracted individuals for which scRNA data was also available, leaving 527 individuals (N=171 AD cases and N=184 healthy controls and N=172 specified as O (other)) for analyses. For these individuals, we further selected variants known to associate with AD from previous GWAS^29,30^. Only variants for which all three genotypes were present in at least 5% of the total population were tested. Quality control of genotype data was performed with PLINK (v1.90b4.6 and v2.00a2.3)^50,51^. Liftover of the genetics data was performed with liftOver R-package (v1.10).

### Bulk AML RNAseq data

The bulk AML RNAseq datast was obtained from Severens, et al^43^. This dataset consisted of 3,656 individuals and 60,660 transcripts. First, individuals having ≤ 20,000 or ≥35,000 zero measurements were removed. Next, genes that were measured in less than 90% of the individuals were removed. The resulting matrix (23,418 genes x 3,383 individuals) was normalized using median ratio normalization^52^. The dataset was comprised of five different source datasets, as such, batch correction was done using Combat from the R-package sva(v 3.46.0)^53^. BiomaRt^54^ was used to translate the ensembl gene IDs to HGNC gene symbols.

### Cell type annotation

The DLPFC2 dataset was used to identify marker genes as it was already annotated. For excitatory neurons (n = 3.154), inhibitory neurons (n = 457), astrocytes (n = 456), oligodendrocytes(n = 283), microglia (646), OPCs (n = 274) and endothelial cells (n = 517) markers were identified. First, pseudo bulk data was generated for each cell type and concatenated, resulting in a gene by individual matrix, in which each individual is present seven times (one for each cell type). Then, for each cell type a differential expression analyses, using Wilcoxon-rank sum test, was performed where the groups were defined by whether the measurement was from the respective cell type (group 1) or not (group 2). A gene was considered a marker gene when P ≤ 5 × 10^-10^ and log_2_ fold-change ≥ 3. Next, with these markers the cells from the TREM, BA10 and BA9 datasets were annotated. This was done using the AddModuleScore function from Seurat^55^, which assigns a score to every cell for each cell type based on the expression of cell type markers. Each cell was annotated as the cell type for which the score was the highest. However, if the second highest score was within 25% of the highest score, it was annotated as hybrid and removed for subsequent analyses.

### Generating pseudo bulk data

For each dataset, for each cell type, pseudo bulk was generated. Aggregation was done based on the binary expression pattern, since the percentage of zeros for a gene in a cell population is highly associated with its mean expression^56^ and aggregating based on the percentage of zero results in less false positive in downstream analyses opposed to aggregating based on the mean^57^. Next, for each cell type, all the datasets were combined, and genes expressed in less than 10% of the individuals were removed. Then, the expression was normalized using median ratio normalization^58^ and batch correction was performed using ComBat from the sva R-package (v3.46.0)^53^. The DLPFC2 dataset consisted of 60 batches and the other datasets were each considered a batch, as such, in total there were 64 batches. Batch correction was confirmed with a PCA and visually inspecting the principal components and visually inspecting boxplots of the individuals’ gene expressions. For the endothelial cells only the DLPFC2 dataset was used.

### Shared variability representation of gene sets

Starting with a gene-by-sample expression matrix we first subset the genes that belong to a specific gene set. Using the subsetted gene-by-sample matrix we first scale each gene such that the mean = 0 and the standard deviation = 1, then we perform a principal component analysis using the prcomp function from the stats R-package. If the first principal component explains ≥10% of the total variance of the gene set, and the gene set is comprised of at least 5 genes, then we store the principal component in the new gene set-by-sample matrix (**Supp. Fig 2**).

### Gene sets

We used four different gene sets. Three of these gene set databases (KEGG^33^, TRRUST Transcription Factors^42^ and Metabolomics Workbench^32^) we downloaded from the webserver of enrichR^59^: https://maayanlab.cloud/Enrichr/#libraries. The microRNA database was downloaded from miRTarBase^31^, where we only considered functionally validated microRNA targets.

### QTL analyses

QTL identification was performed using a linear model which was defined as follows:

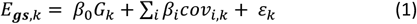

where *E*_*gs*_ is the shared variable representation of gene set *gs* (when performing gsQTL), or, alternatively, the gene expression of a single gene (when performing eQTL), for an individual *k. G*_*k*_ is the genotype dosage of an individuals of the SNP of study (0-2) . *Cov*_*i*,*k*_ are the covariates for each of the individuals. *ε*_*k*_ is an error term that gets minimized. In both, the GSQTL and eQTL analyses, we adjusted for age, sex, diagnosis, dataset, and the first five gene expression PCs. For the bulk AML analyses, *G* represents subtype assignment (0-1) and we adjusted for sex and tissue of origin (bone marrow and peripheral blood).

### Transcription factor validation experiment

Using the healthy individuals and the excitatory neuron dataset, we evaluated the association between the expression of transcription factors (TFs) and shared representation derived from the genes targeted by the respective TF. Targeted genes were derived from the trust transcription factor database^42^. We only evaluated TFs that were present in both the trust database as well as in our expression data. Potential covariance caused by effects of age, sex, dataset and Braak stage on the gene expression were regressed out. We only considered TF targeted gene sets, when there were minimally five target genes, and the shared variability had to explain at least 5% of the total variance of the gene set. The association between the expression of the TF and the expression in the shared variable representation was evaluated using Spearman’s rank correlation coefficient. The resulting p-values were adjusted for testing multiple TF target gene tests using the Benjamini-Hochberg Procedure, and significance of the association was assumed at P_FDR_ ≤ 0.05.

### ORA: Over representation analysis

Overrepresentation analysis (ORA) was performed using the fisher exact test for each variant separately including the genes that are nominally significant, i.e. eQTLs for which P ≤ 0.05. The Benjamini-Hochberg Procedure was used to corrects for testing multiple gene sets, and a gene set was assumed significant at P_FDR_ ≤ 0.05.

### GSEA: Gene set enrichment analysis

Gene set enrichment analysis (GSEA) was performed using the R-package fgsea^41^ (v 1.24.0). For each variant, the βs of the associations with the genes were used to rank the genes, that served as input for the gene set enrichment analysis. The resulting p-values were adjusted for testing multiple tests per variant using the Benjamini-Hochberg Procedure, and significance of the association was assumed at P_FDR_ ≤ 0.05.

