## Supplementary Figures for "gsQTL: Associating genetic risk variants with gene sets by exploiting their shared variability"

Gerard A. Bouland[1,2,], Niccolo Tesi[1,3] , Ahmed Mahfouz[1,2,*] and Marcel J.T. Reinders[1,2,*]

[1] Delft Bioinformatics Lab, Delft University of Technology, Delft, The Netherlands

[2] Department of Human Genetics, Leiden University Medical Center, Leiden 2333ZC, The Netherlands

[3] Section Genomics of Neurodegenerative Diseases and Aging, Department of Clinical Genetics, Vrije Universiteit Amsterdam, Amsterdam UMC, Amsterdam, The Netherlands

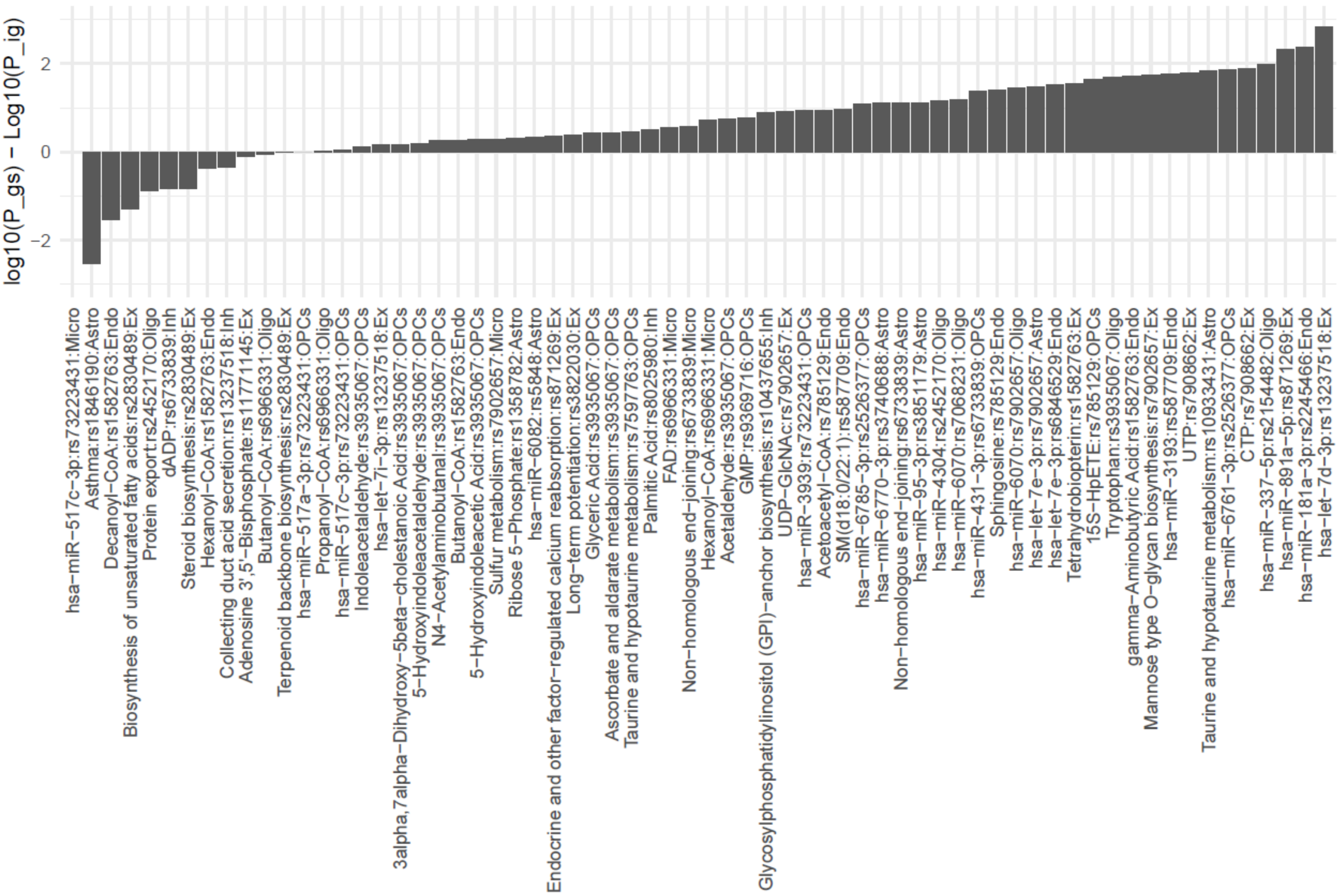


**Supplementary Figure 1:** A bar plot where the x-axis represent significantly identified gsQTLs and the y-axis represents the difference between the -log10(p-value) of the gsQTL and the -log10(p-value) of the most significant individual gene that was part of the respective gene set used to calculate the proxy values

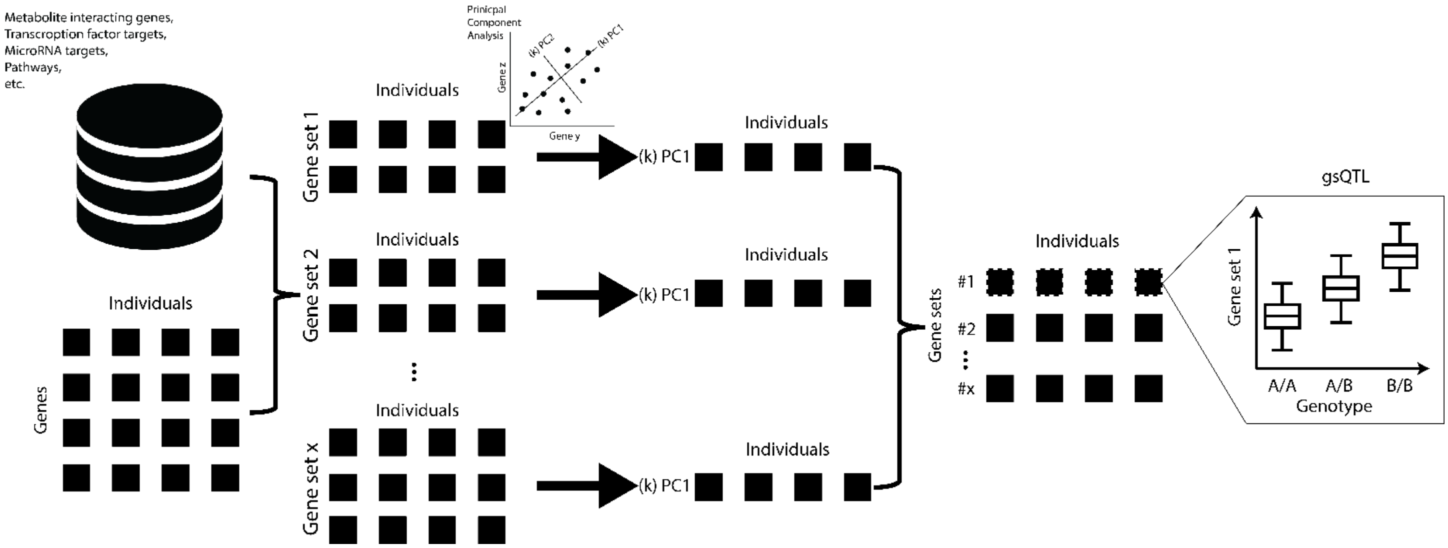


**Supplementary Figure 2:** Visual representation of the approach. First, we take as input a gene expression matrix where the rows represent the genes and the columns represent the individuals or samples. Then, for the selected gene sets (e.g. pathways, metabolite interacting genes, transcription factor or microRNA targets), the respective genes are subsetted from the input dataset. For each gene set PCA is performed, either a linear PCA or a kernel-PCA. Next, PCs are filtered based on the number of input genes and based on percentage variance explained. Finally, QTL analyses are performed on the PCs.

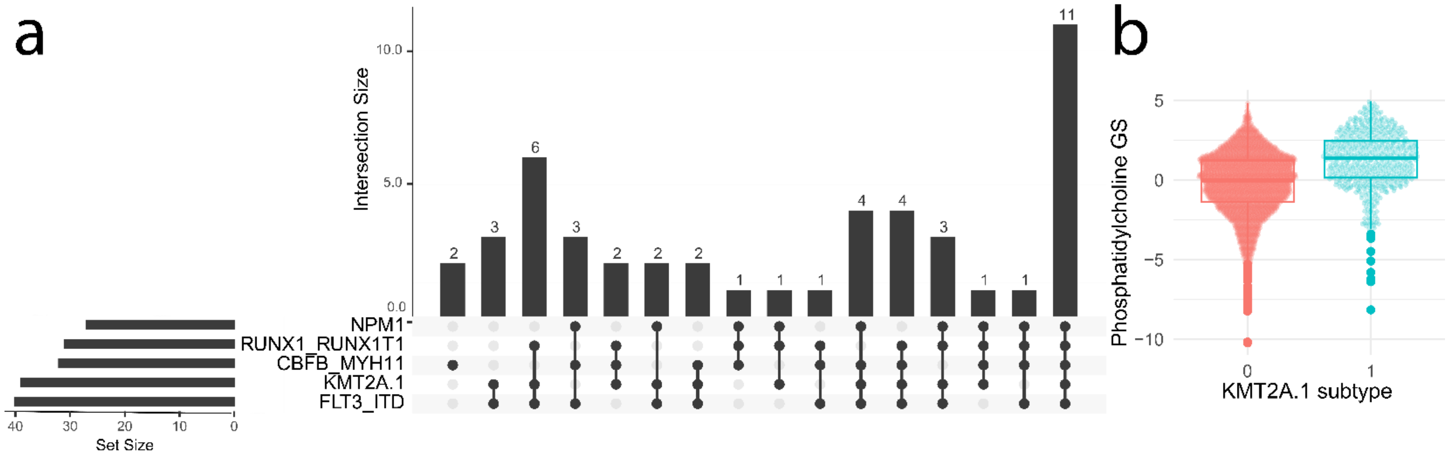


**Supplementary Figure 3: A)** Upset plot of the overlap of significantly identified proxy metabolites between the AML subtypes. **B)** Boxplot of the phosphatidylcholine GS-metabolite. X-axis represents KMT2A.1 status where 0 is negative and 1 is positive. Y-axis represents the expression value.
